# Co-contraction uses dual control of agonist-antagonist muscles to improve motor performance

**DOI:** 10.1101/2020.03.16.993527

**Authors:** Christopher M. Saliba, Michael J. Rainbow, W. Scott Selbie, Kevin J. Deluzio, Stephen H. Scott

## Abstract

Co-contraction of agonist-antagonist muscles is commonly observed when performing difficult motor tasks. The benefit of co-contraction is thought to be zero-delay corrections to unexpected disturbances from increased intrinsic muscle impedance. We used upper-limb postural and tracking tasks to characterize the effects of co-contraction on motor corrections to loads applied to the limb. We systematically controlled pre-perturbation muscle activity and showed that co-contraction improves subsequent corrective responses in both tasks. However, substantial improvements in the corrective response are only observed at the time when neural feedback pathways can also contribute. We demonstrate that muscle impedance appears to play a minor role in improving performance. Instead, co-contraction engages a dual agonist-antagonist control strategy to counter disturbances, that is distinct from the control strategy used when not co-contracting or selectively pre-activating a single muscle group. Critically, we showed that this dual agonist-antagonist control strategy improved performance even at low levels of co-contraction.

## Introduction

A common strategy to perform difficult motor tasks is to co-activate antagonist muscles. For example, we increase activity of muscles in our arm to increase grip and downward force to turn a screw. Increases in co-contraction have been observed experimentally by simply asking participants to maintain their forearm in an upright posture with varying amounts of weight held in their hand^1^. As the weight increases, the level of co-contraction also increases in the elbow flexors and extensors (See also ^2^). As well, learning to reach with novel loads leads to an initial strategy to co-contract limb muscles^3–6^, and increased co-contraction of leg muscles is observed when anticipating a slippery surface during gait^7^. Why co-contract when it increases the required effort (i.e. metabolic cost) without any increase in net force output?

The most common explanation for co-contracting agonist and antagonist muscles is for impedance control^8^ by increasing muscle stiffness and damping^1^. Increasing muscle impedance through cocontraction has been suggested to provide an immediate response to unexpected disturbances without the problem of delays for neural control^9,10^. Thus, co-contraction improves stability between the body and the environment by exploiting the intrinsic properties of muscle.

However, conventional techniques to measure muscle or limb impedance include the contribution of neural feedback processes. Direct measures of limb stiffness (change in length for a given change in force) quantify the change in length (or hand position) when a load is applied to the limb. Notably limb stiffness is quantified >200 ms after the application of a load^11–16^, and thus, the response to the load includes the spinal, short-latency feedback response starting at ~25ms, and the long-latency cortical feedback response starting at ~60 ms. Therefore, estimates of muscle stiffness include contributions from both muscle and neural feedback^17,18^. While one could try to estimate stiffness immediately after a disturbance, before spinal responses influence force output (~50 ms), motion in this time period is dominated by limb inertia making it difficult to estimate stiffness or damping^19^.

The purpose of the present study was to understand the influence of co-contraction on motor function. Rather than eliciting co-contraction through destabilizing dynamics, we systematically controlled the level of co-contraction during the task prior to the perturbation. Using this paradigm, we first confirm that co-contraction improved corrective responses during postural control and endpoint tracking tasks. Systematic increases in activity of only one muscle group provided far less improvement in performance than co-contraction. Faster corrective responses associated with co-contraction were generated by the combined effects of neural feedback responses and mechanical impedance. Critically, a key benefit of co-contraction appears to be that both stretched and shortened muscles remain active after the perturbation and contribute to feedback corrective responses. In contrast, feedback corrections are limited to the muscle group that was pre-activated whether it is stretched or shortened. We demonstrated the benefit of this dual agonist-antagonist control strategy using a computational model.

## Results

### Performance improves with co-contraction during a postural task

In our first experiment, participants performed a postural perturbation task using a Kinarm Exoskeleton robot^20^ (Kinarm, Kingston, Ontario, Canada). They began each trial by moving their finger to a central target. A perturbation torque-pulse flexed or extended the elbow, displacing the fingertip from the target. Participants completed the task successfully if they returned their fingertip to the target in <500 ms and held it in the target for >1000 ms. The task was performed with background torques of 1-5 Nm applied at the elbow to pre-activate either the elbow extensors or flexors (Figure 1a). The preactivated muscles are considered here to be the agonists. Participants also performed the task with no background load. In this case, the subject started each trial by co-contracting the elbow flexor and extensor muscles to activity levels corresponding to 1, 3, or 5 Nm background torques using visual feedback (Figure 1b).

**Figure 1:**
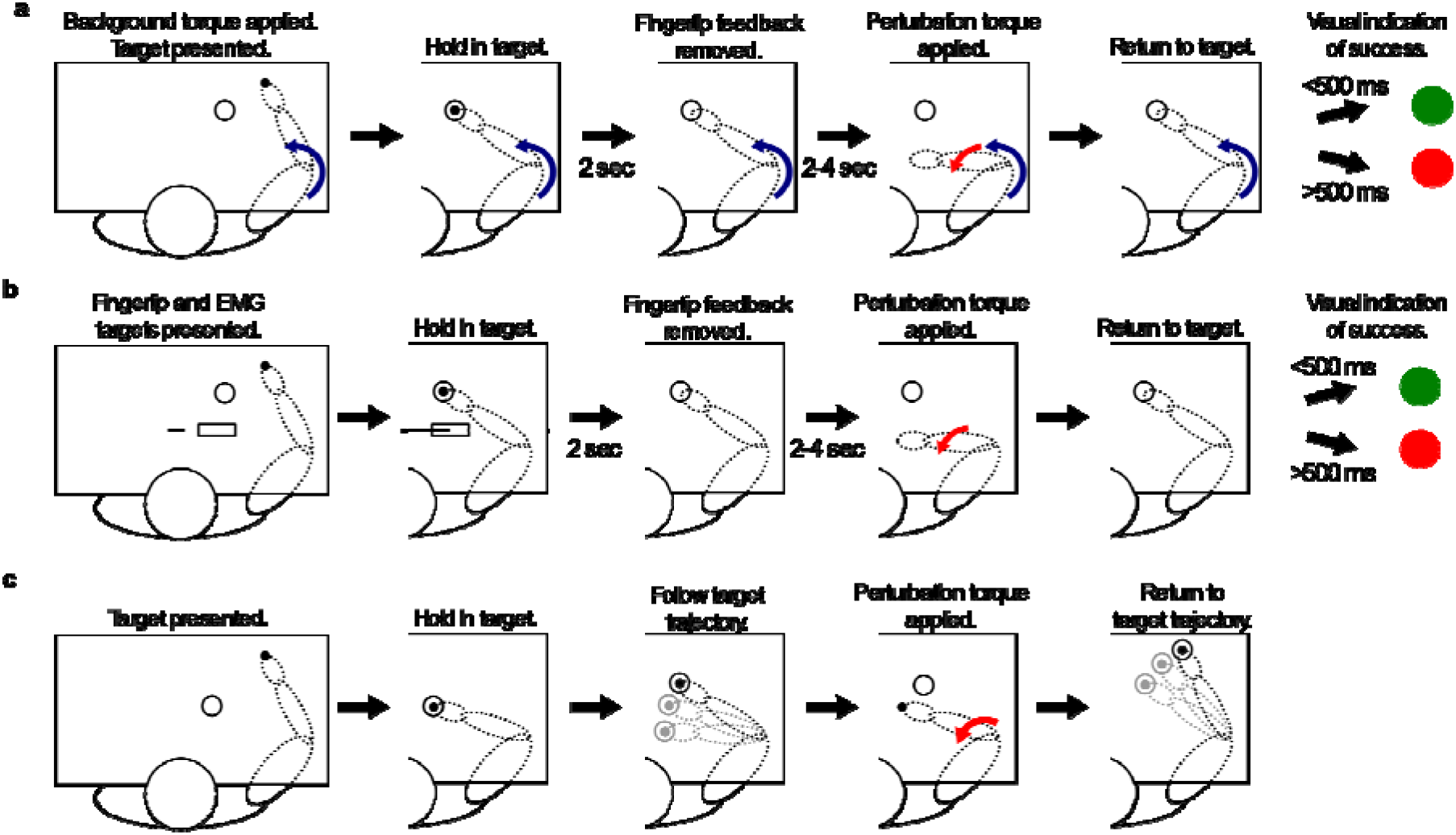
(a) Postural perturbation task with a background load that pre-activates the elbow extensors or flexors. Participants must keep their fingertip in a stationary target while resisting the background load. A perturbation torque pulse is applied, and participants must return their fingertip to the target as quickly as possible. (b) Postural perturbation task with targeted co-contraction of the elbow flexors and extensors. Participants must co-contract to a desired level of muscle activity using visual feedback while holding their fingertip in a stationary target. A perturbation torque pulse is applied, and participants must return their fingertip to the target as quickly as possible. (c) Target tracking task. Participants must track a moving target with their fingertip. A perturbation torque pulse is applied, and participants must return their fingertip to the moving target as fast as possible. Trials are performed with and without active co-contraction of the elbow flexors and extensors.

We found a clear improvement in the performance of participants when they co-contracted prior to the postural perturbation as compared to when they countered background loads (Figure 2a). Load condition and level of muscle activity had a significant effect on task success rate with no significant interaction (F(condition) = 25, p = 2.1e-6; F(level) = 4.2, p = 0.028; F(condition x level) = 2.3, p = 0.071). Task success was 56% when at the lowest level of co-contraction and increased to 71% at the highest level (p = 0.021, paired t-test between participants). In contrast, task success was only ~35% when the muscle group pre-activated by the background loads (agonist muscle group) was stretched or shortened by the perturbation. There was a small increase in success to 45% for the agonists stretched trials at the highest level of background load. Increases in performance when co-contracting compared to countering background loads (an additional 23% to 39% of the total trials completed successfully) were statistically significant (p = 0.013 for co-contraction vs agonists stretched at low muscle activity, p < 0.01 for co-contraction vs all others, paired t-tests between participants).

**Figure 2:**
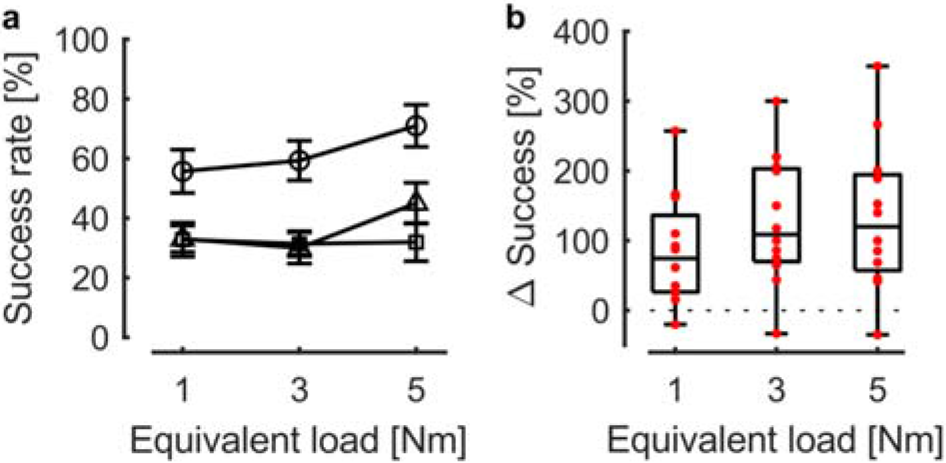
(a) Task success rate for the postural perturbation task across all subjects for the three experimental loading conditions: agonist stretch (triangle), agonist shorten (square), and co-contraction (circle). (b) Relative changes in task success for the postural perturbation task with co-contraction compared to pre-activation of a single muscle group. Red dots indicate individual subjects. Equivalent load indicates the level of background muscle activity from resisting background loads or co-contracting.

Eleven of the twelve participants improved performance at all three levels of muscle activity when co-contracting as compared to when the agonists were stretched or shortened (Figure 2b). Relative increases in performance of up to 350% were achieved, with a median performance improvement of ~100%.

The pattern of corrective responses when participants co-contracted was clearly different as compared to when only one muscle group was pre-activated (Figure 3). Background muscle activity level had a significant effect on peak displacement with an additional interaction effect from loading condition (F(condition) = 2.8, p = 0.083; F(level) = 155, p = 1.1e-13; F(condition x level) = 20, p = 2.5e-9). Loading condition, activity level, and their interaction had significant effects on the return time following the perturbation (F(condition) = 26, p = 1.7e-6; F(level) = 28, p = 9.3e-7; F(condition x level) = 3.1, p = 0.026).

**Figure 3:**
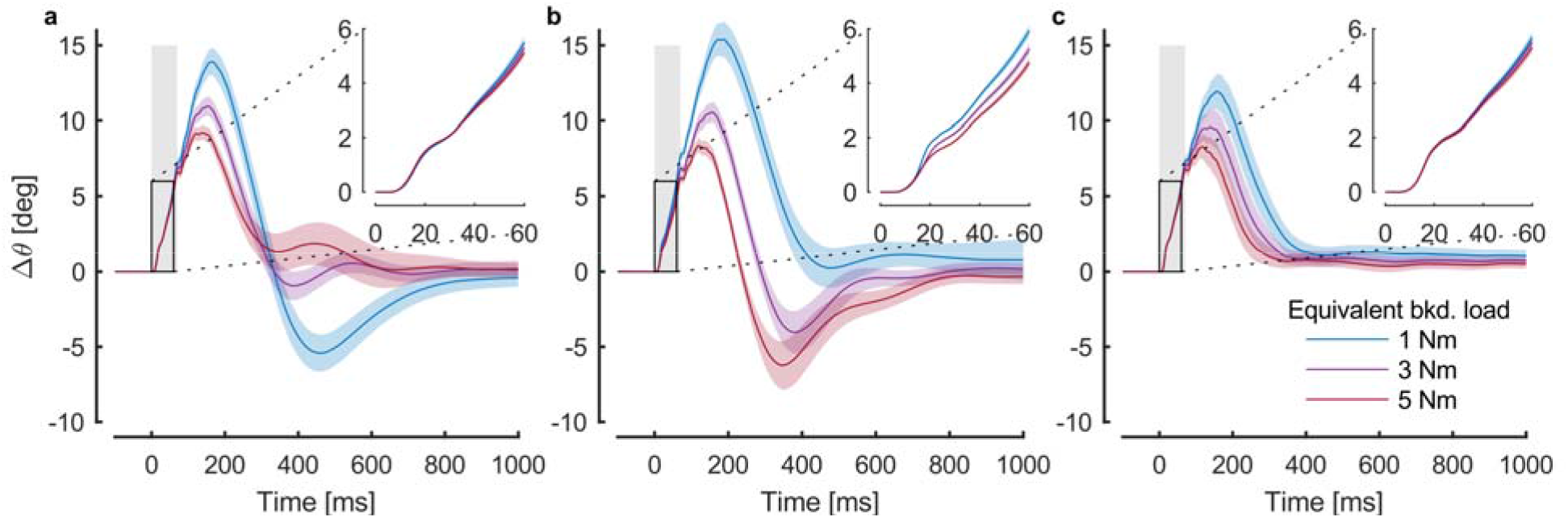
Change in elbow angle following mechanical perturbations for the three experimental loading conditions: (a) agonist stretch, (b) agonist shorten, and (c) co-contraction. Background muscle activity level is indicated by line color and perturbation onset is at 0 ms (grey box). Group results (mean and standard error) are shown.

Detailed analysis of the changes in elbow motion highlights several critical differences in limb motion following perturbations. First, there were substantial differences in the pattern of elbow motion even at the lowest load level whether the agonists were stretched, the agonists were shortened, or all muscles were co-contracted (blue lines in Figure 3). Maximal displacement was similar when only one muscle group was pre-activated, whereas when they were co-contracted it was 14% (2.0 degrees) smaller than when the agonists were stretched (p = 0.21, paired t-test between participants) and 23% (3.5 degrees) smaller than when the agonists were shortened (p = 0.0027, paired t-test between participants). There was a substantial overshoot when the agonists were stretched that does not occur when the agonists were shortened, or when both flexor and extensor muscle groups were co-contracted. Although the elbow returned to the target angle at about the same time for the latter two conditions, there was more variability when the agonists were shortened by the perturbation (standard deviation 1.3 degrees) than when co-contracting (standard deviation 0.5 degrees) and this likely leads to greater success for cocontraction trials (p = 1.2e-4, two sample F-test using elbow angle 500 ms post-perturbation).

A second important kinematic feature was the small change in the initial post-perturbation elbow motion for different background loads or co-contraction levels (Figure 3 insets). The largest difference was observed when the agonists were shortened, resulting in a 21% decrease in limb motion at 50 ms. However, the difference in motion during co-contraction was only 5%. This suggests that intrinsic muscle properties influence initial limb motion but the effect at this short time-scale is limited.

Third, increases in background load resulted in a clear drop in peak displacement. Peak displacement decreased by 30 to 45% from the lowest to highest background activity levels, depending on the load condition (p < 0.01, paired t-tests between participants). At the highest load levels, the maximum displacement was similar whether the agonists were stretched, the agonists were shortened, or both muscle groups were co-contracted.

Fourth, the most critical difference across the load conditions was related to the presence or absence of overshoot at the end of the corrective response. The overshoot decreased as the background load increased when the agonists were stretched, whereas it increased when the agonists were shortened. Increases in co-contraction resulted in a slightly faster return to the target angle (200 ms decrease in return time between low and high levels of muscle activity, p = 0.019) while continuing to have minimal overshoot. Thus, co-contraction resulted in a simple stereotyped pattern of motor correction, whereas pre-activating only one muscle group resulted in more complexity in the motor correction dependent on the level of background muscle activity.

A critical difference between co-contraction and the other load conditions was the patterns of muscle activity during the motor corrections (Figure 4). When the agonist muscles were stretched, they responded with a burst of activity during the long-latency epoch. As background agonist activity increased, an earlier burst of activity, during the short-latency epoch, also occurred. The magnitude of this burst increased with the level of background muscle activity, commonly called gain scaling^21–25^. Importantly, at all levels of background activity, there was effectively no motor response in the antagonist muscles.

**Figure 4:**
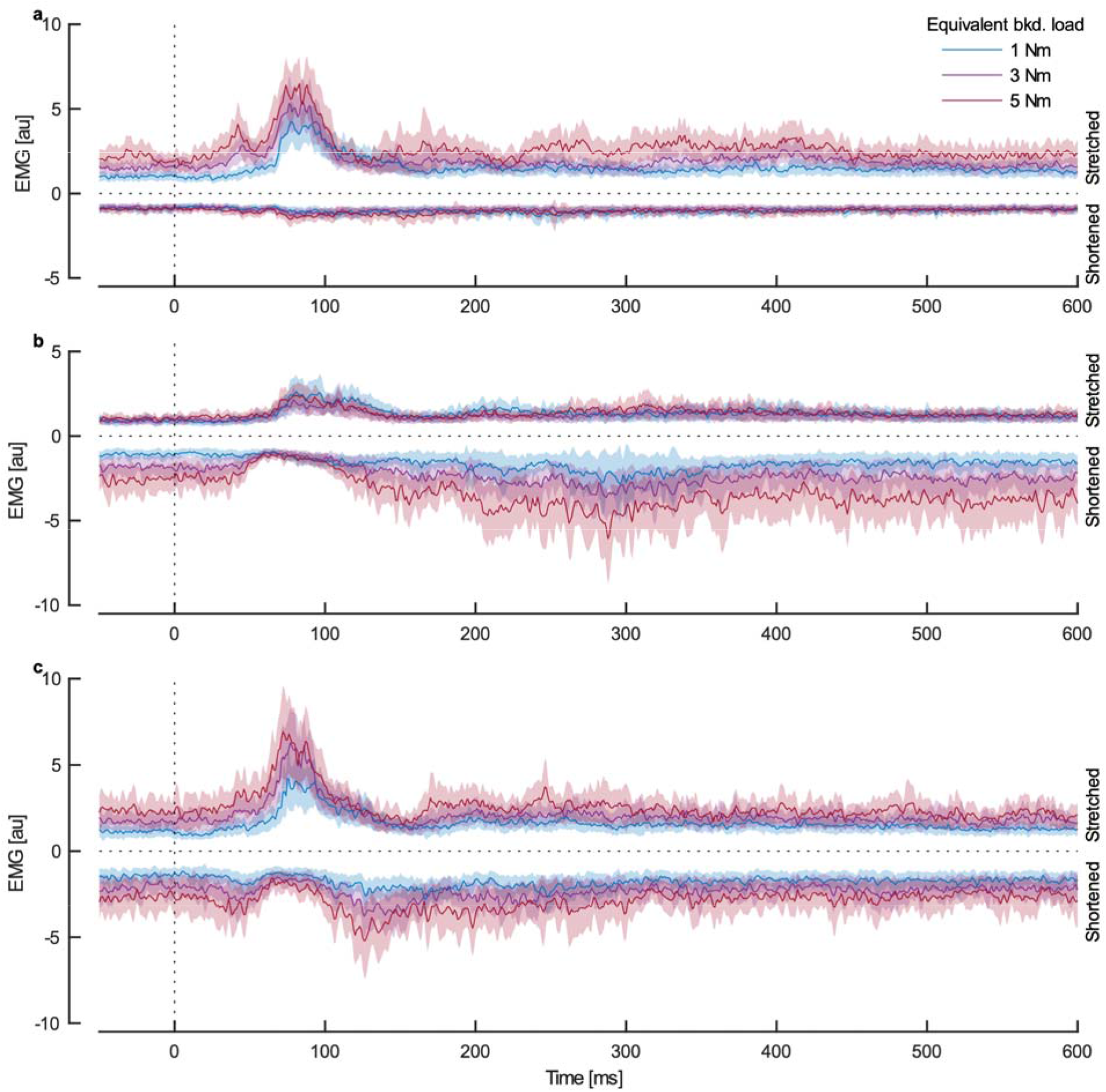
Activity of the triceps lateral head and biceps following mechanical perturbations that flex the elbow. (a) Stretched triceps pre-activated by background load (agonist stretch). (b) Shortened biceps pre-activated by background load (agonist shorten). (c) Muscles co-contracted. Increases in activity of the stretched triceps are plotted in the positive direction and increases in activity of the shortened biceps are plotted in the negative direction. Background muscle activity level is indicated by line color and perturbation onset is at 0 ms. Group results (mean and standard error) are shown. See Supplementary Material for additional muscle activity figures.

When the agonist muscles were shortened, they were inhibited and briefly dropped to minimal activity for all background load levels. Activity increased again from 120 ms to 350 ms post-perturbation. This increase scaled with the level of background agonist activity. At all levels of background agonist activities, the antagonist muscles exhibited minimal response to the perturbation.

Finally, when co-contracting, the corrective response engaged both the stretched and shortened muscles. The response of the stretched muscles was qualitatively the same as observed during the agonists stretched condition. The shortened muscles responded with inhibition during the long-latency epoch that scaled with background activity. There was a sudden increase in activity following the inhibition that occurred between 90 ms and 170 ms post-perturbation. This burst scaled with background activity and was initiated prior to hand reversal, likely preventing overshoot when returning to the target^26^. Additionally there appears to be a small burst of activity during the short-latency epoch in the shortened muscles, suggesting that muscles may co-contract reflexively following the perturbation^27^. However, this reflexive co-contraction was not present in all subjects or muscles.

### Model to observe contributions of impedance and feedback control

We developed a model that captures two important aspects of co-contraction: increased mechanical impedance and feedback control of both stretched and shortened muscles (dual agonist-antagonist control). The model consists of a rigid link with a pin joint at one end to represent the forearm, hand, and elbow. Two torque actuators act as the elbow extensors and flexors. These actuators act in opposite directions, can only generate pulling torques, and are governed by a first order approximation of activation dynamics^28^. The actuators are driven by an optimal feedback controller designed to return the link to the starting posture following a pulse perturbation. Co-contraction is added to the model in two ways. First, the stiffness and damping at the pin joint are increased at higher levels of co-contraction based on the force-length and force-velocity properties of muscle^28,29^. Second, the baseline activity of the actuators is increased, allowing increases as well as decreases to zero activity to counter the perturbation.

We wanted to examine the individual contributions of increased mechanical impedance and increased background activity associated with co-contraction. The influence of impedance without feedback control was captured by the passive simulations (Figure 5a). Increasing impedance reduced the maximum displacement, the overshoot when returning to the target, and the return time of the system. Simulations with feedback control and baseline activity of agonist and antagonist muscles while muscle impedance was kept fixed, isolated the effect of the dual agonist-antagonist control strategy (Figure 5b). Dual control greatly reduced the maximum displacement of the system and eliminated target overshoot even without increases in impedance. Importantly, good performance occurred even at the lowest level of co-contraction. Finally, when impedance was increased with baseline activity (Figure 5c), performance slightly improved providing a slightly faster return time.

**Figure 5:**
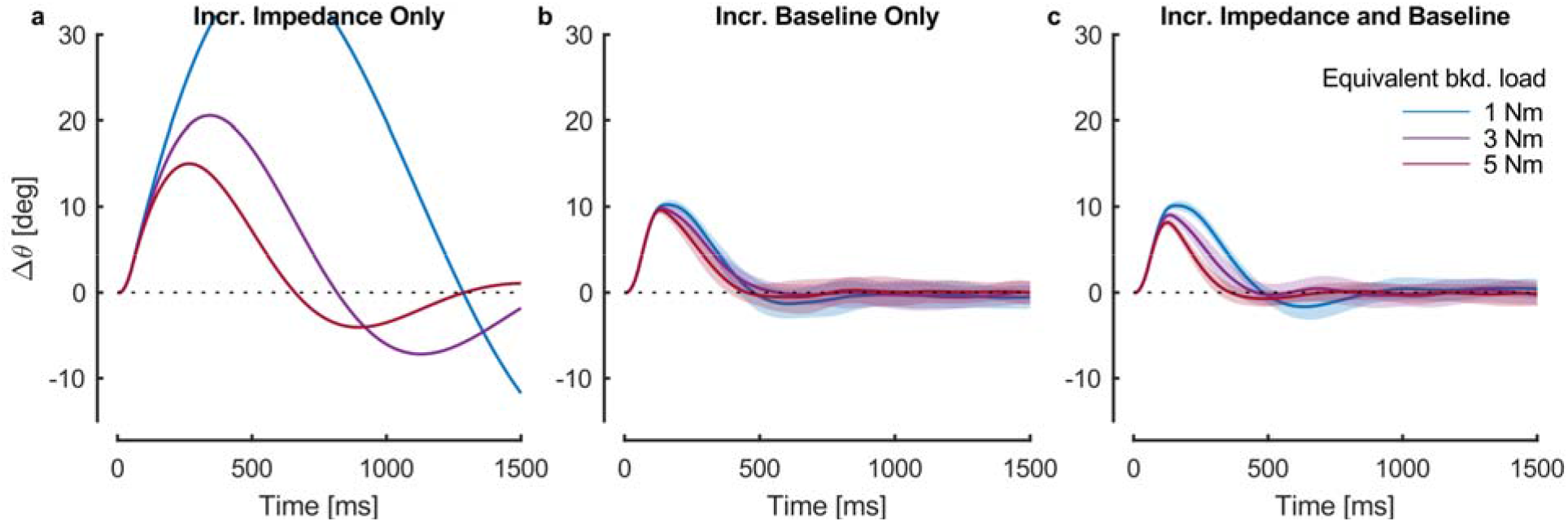
Simulation results (change in elbow angle following mechanical perturbations) showing the effects of muscle intrinsic impedance and feedback control. (a) Passive simulations in which impedance increases with co-contraction, but there is no feedback control. Note that the trajectory of the response at the lowest level of impedance extends beyond the limits of the axes in order to maintain a constant scale across the plots. (b) Simulations in which there is feedback control and baseline activity increases with co-contraction, but impedance remains at the lowest level. (c) Simulations with both feedback control and increasing impedance with co-contraction. Line color represents the equivalent muscle activity level used to set baseline activity and/or impedance. Mean and standard deviation of 20 simulations are shown.

We wanted to verify the benefits of having two muscle groups active for control as compared to when only one muscle group was active. In each case, the controller was optimized assuming that both actuators contribute to the feedback response. Simulations in which only the stretched (Figure 6ad) or shortened (Figure 6be) actuator could respond to the perturbation and simulations in which both could respond (Figure 6cf) represent the agonists stretched, agonists shortened, and co-contraction experimental conditions, respectively. In all three cases, the impedance increased in accordance with the level of baseline activity. The results of these simulations share many key characteristics with the experimental results.

**Figure 6:**
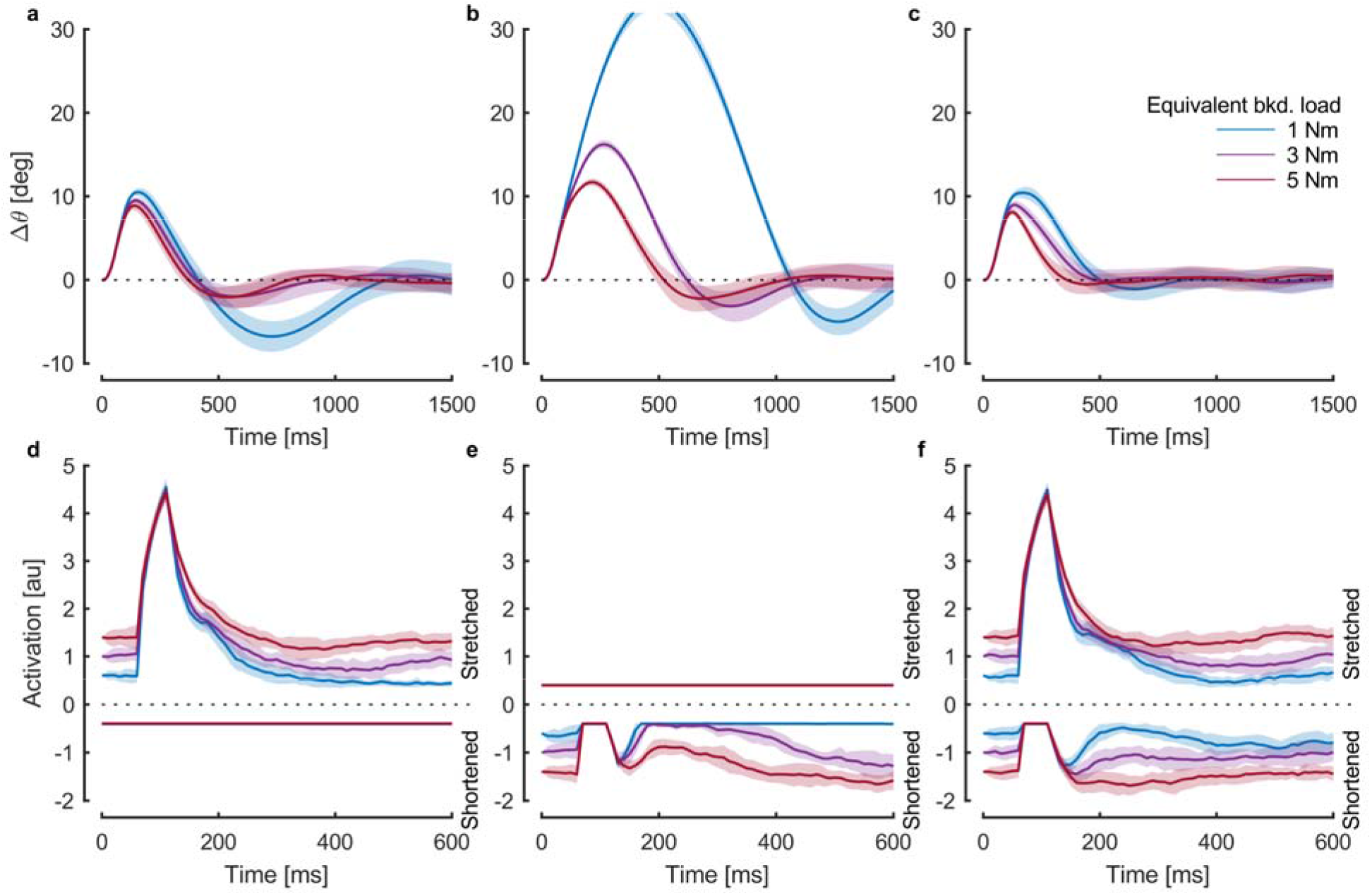
Change in elbow angle following mechanical perturbations from simulations where (a) only the stretched actuator responds, (b) only the shortened actuator responds, and (c) both actuators respond. The corresponding actuator activities are shown in (d-f). Increases in stretched actuator activation are plotted in the positive direction and increases in shortened actuator activation are plotted in the negative direction. Line color represents the equivalent muscle activity level used to set baseline activity and impedance. Mean and standard deviation of 20 simulations are shown.

The most important result is that control with both actuators (dual agonist-antagonist control) eliminated the overshoot and oscillations that occurred when only one actuator was active. Further, increasing background activity reduced maximum displacement in all conditions, and reduced overshoot when the single active actuator was stretched.

The patterns of actuator activity in the simulations were qualitatively similar to the patterns of muscle activity used by the participants (Figure 6d-f). When pre-activated, the stretched actuator responded with a burst of activity followed by suppression. At the lowest baseline level, this suppression was clipped due to the restriction that the actuator could only generate torque in one direction. When the shortened actuator was pre-activated, it responded with inhibition that was clipped at zero activity for the lower baseline levels. When both actuators were pre-activated, their responses followed the same patterns as when individually pre-activated. However, with both active, clipping was less common and principally only observed briefly in the shortened actuator at the lowest baseline level. While the model captures many features of human motor corrections, it should be noted that unlike in the experiment, the magnitude of the burst of the stretched actuator did not increase with baseline activation. Additionally, overshoot did not increase with baseline activation when only the shortened actuator responded.

### Performance improves with co-contraction during a tracking task

We used a tracking perturbation task to test if co-contraction also improved performance during movement. In this task, participants began each trial by moving their fingertip to a stationary target (Figure 1c). The target began to move at a constant speed along a trajectory that required only elbow flexion or extension. Once the end of the trajectory was reached, the target reversed direction and followed the same trajectory back to the starting point. At a random time, when the target was near the center of the trajectory, a perturbation torque pulse (5 Nm, 50 ms) flexed or extended the elbow, displacing the fingertip from the target. This created four perturbation conditions: extension movement with flexion perturbation, extension movement with extension perturbation, flexion movement with extension perturbation, and flexion movement with flexion perturbation. Participants were instructed to perform the task with and without active antagonistic co-contraction of the muscles spanning the elbow.

We found a clear improvement in the performance of participants when they co-contracted (Figure 7). Co-contracting reduced deviations from the desired elbow motion caused by the perturbation regardless of the movement and perturbation directions and eliminated overshoot at the end of the corrective response. There was a modest decrease (12%) in the initial (50ms) post-perturbation elbow motion when co-contracting.

**Figure 7:**
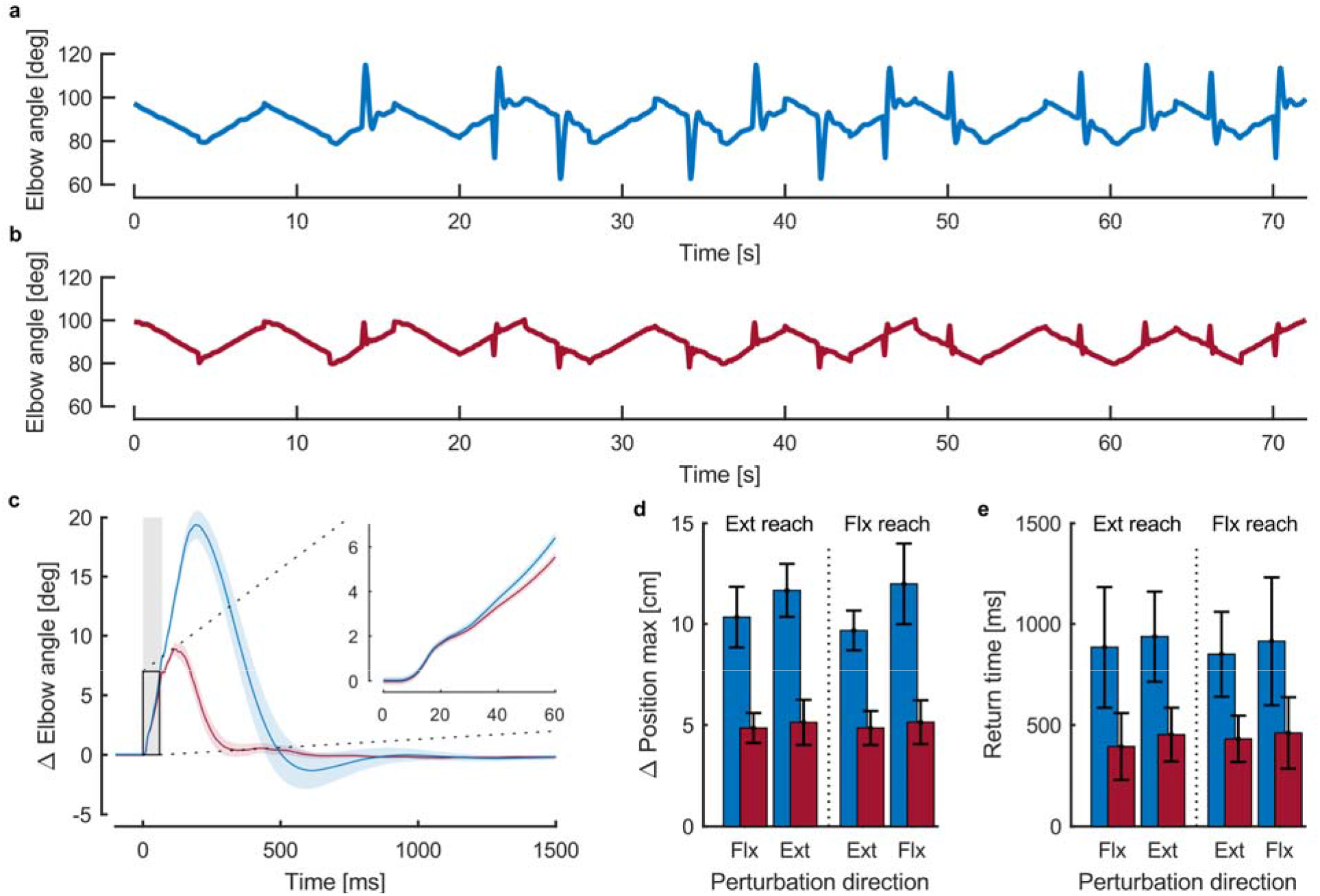
Elbow angles during repeated out and back tracking movements subject to mechanical perturbations by an exemplar subject (a) without and (b) with active co-contraction. (c) Elbow angle deviation from the target trajectory following mechanical perturbations without (blue) and with (red) active co-contraction. Perturbation onset is at 0 ms (grey box). Group results (mean and standard error) are shown. (d) Maximum distance from the fingertip to the target and (e) time to return to the target following mechanical perturbations without (blue) and with (red) active co-contraction. Group results (mean and standard deviation) are shown.

Co-contraction, perturbation condition, and their interaction had a significant effect on the maximum displacement of the fingertip (F(co-contraction) = 287, p = 3.9e-8; F(condition) = 24, p = 8.1e-8; F(co-contraction x condition) = 21, p 2.6e-7). Return time was only significantly affected by co-contraction (F(co-contraction) = 81, p = 8.4e-6; F(condition) = 0.53, p = 0.66; F(co-contraction x condition) = 0.17, p = 0.92). Maximum displacement and return time both decreased between 50% and 57% when co-contracting, depending on the perturbation condition (p < 0.001 for all conditions, paired t-tests between participants).

When performing the task without co-contracting, the perturbation response was limited almost entirely to the stretched muscle, that generated a burst of activity during the long-latency epoch (Figure 8, blue traces). The shortened muscle generated little to no response. In contrast, when co-contracting the motor response involved both the shortened and stretched muscles (Figure 8, red traces). The stretched muscle generated a greater short and long-latency response than in the no co-contraction condition. In addition, the shortened muscle produced inhibition during the long-latency epoch followed by a burst of activity starting at approximately 100 ms post-perturbation that lasted between 100 ms and 200 ms. As in the static task, this burst occurred prior to hand reversal, likely preventing overshoot in the return to the target.

**Figure 8:**
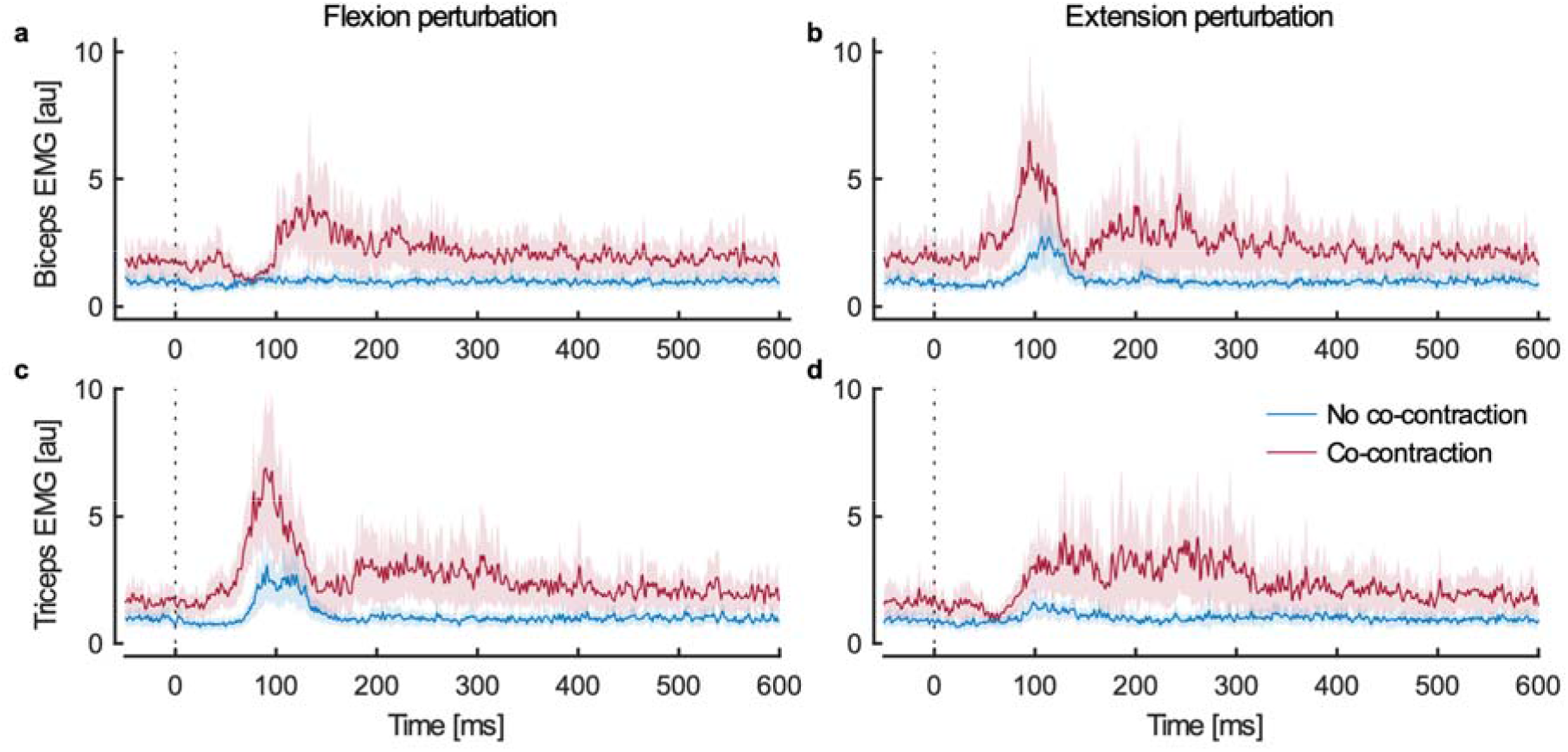
Activity of the biceps (top row) and triceps (bottom row) during the target tracking task with flexion (first column) and extension (second column) perturbations. Group results (mean and standard error) are shown for trials without (blue) and with (red) active co-contraction. Perturbation onset is at 0 ms.

## Discussion

We show in the present study that co-contraction of agonist and antagonist muscle groups improves the success of corrective responses to unexpected disturbances during both postural control and tracking tasks. We systematically controlled the level of co-contraction participants generated prior to the perturbation, rather than allowing them to select the level of co-contraction^3,30,31^ □. In doing so, we could isolate mechanisms and patterns of the neuromuscular response used when co-contracting.

The key benefit of co-contraction has been attributed to an increase in muscle impedance generating zero-delay responses to help counter mechanical disturbances^10,32^. However, our results highlight that initial limb motion is not substantively impacted by changes in the level of muscle activity (see also^19^). While changes in impedance may not impact the immediate response of the limb, impedance does change the corrective response over a longer period of time. These effects are demonstrated by the passive behavior of our model. With low impedance, the system took longer to recover from the perturbation and the response included large overshoot and oscillations. Increasing muscle impedance produced faster responses and greatly reduced overshoot and oscillations. Thus, impedance does improve the perturbation response but the effects are not noticeable until later in the response when neural feedback is also active (>50 ms).

A possible limitation of our model is that we do not include short-range muscle stiffness^33–35^ and thus may underestimate the effects of muscle impedance. We chose not to include short-range stiffness for two reasons. First, the perturbation torque stretched the elbow extensors and flexors beyond the shortrange, so the majority of the response does not have active short-range stiffness^19^. Second, we observed the same benefits of increasing muscle activity through co-contraction during the tracking task when short-range stiffness was not present. While we used an optimal feedback control in our simulations with a 60 ms sensory delay, the results are not specific to this type of control. Other approaches for control with physiologically realistic delays will almost certainly attain similar results.

Of particular note is that rapid corrective responses were generated without overshoot even at the lowest level of co-contraction (equivalent to 1Nm in each muscle group). In contrast, activating a single muscle group more than double that amount resulted in poorer performance. Thus, it is not the total amount of muscle activity (or muscle impedance) that is the key factor when co-contracting, but the fact that both muscle groups are simultaneously pre-activated before the disturbance. Further, the performance improvements observed at this low level of muscle activity show that the benefits of cocontraction can be attained with a low metabolic penalty.

The patterns of EMG during the corrective response highlight that improved performance from cocontraction is principally due to both stretched and shortened muscle groups being actively involved in the corrective response. In contrast, when only one muscle group is pre-activated prior to the disturbance, the other muscle group remains inactive. This failure to engage the non-active muscles appears to reflect a default pattern of reciprocal excitation-inhibition of agonist-antagonist muscles. That is, during voluntary motor actions, excitation of one muscle group tends to be coupled with inhibition of the antagonist muscle group. A key component of this process is likely the Ia inhibitory interneurons in the spinal cord that are part of the disynaptic reciprocal inhibitory pathway^36,37^. When one muscle group is actively recruited during motor actions, these interneurons also receive descending input and synapse onto and inhibit motoneurons of the antagonist muscle group^38–41^. The ongoing inhibition appears to be sufficiently strong that the antagonist cannot be recruited during the feedback response so only the agonist is engaged for control. In addition, the stretch response of muscle afferents in the antagonist may also be diminished when the muscle is actively inhibited due to alpha-gamma coactivation^42^. Performance further degrades at lower levels of agonist pre-activation because muscle activity decreases to zero in the active muscle.

Co-contraction appears to create a fundamental change in spinal circuitry during motor actions. Earlier work found that reciprocal inhibition was significantly reduced at very low levels of cocontraction, and the amount of reduction did not increase at higher levels of co-contraction^39,43^. A key finding was that the reduced reciprocal inhibition could not be explained by the sum of the effects of activating each muscle individually. It was concluded that during co-contraction the Ia inhibitory interneurons were functionally uncoupled from the corresponding motor neurons. This uncoupling could be due to reduced descending drive onto Ia inhibitory interneurons, or the influence of presynaptic inhibition of Ia afferents or Renshaw cell activity onto la inhibitory interneurons^39,44,45^. An additional finding of this earlier work was that the depression of reciprocal inhibition occurred when blocking peripheral feedback to the spinal cord (through induced ischemia) and at the very beginning of dynamic co-contraction (before sensory feedback from the contracting muscles had reached the spinal cord). This indicated that the decoupling of the Ia inhibitory interneurons was the result of a central mechanism specific to co-contraction^43^. Further, another study showed that corticomotoneuronal cells with no inhibitory projections onto antagonist motor neurons were more likely to be active during cocontraction than those with inhibitory projections^39,46^. In combination, these findings suggest that a distinct motor strategy that alters the state of spinal inhibitory circuitry may be used when co-contracting to enable both agonist-antagonist muscles to participate in feedback responses, and correspondingly, improve performance. Importantly, the fact that both agonist-antagonist muscles are used when co-contracting in our tracking task highlights that this is not a simple strategy invoked only for stationary postural control, but a strategy that can also improve performance during movement (at least at the slow speed of our tracking task).

While the hypothesis above focuses on the influence of spinal processes on the neural basis of control, the extent to which supraspinal processes are actively involved during these feedback corrections is unclear. Previous work has highlighted that transcortical feedback pathways are engaged ~60 ms after a perturbation is applied and contribute to feedback responses^17,47^. These studies commonly use step torque loads which necessitate a change in the overall level of muscle activity to counter the applied load. In contrast, the present study used a 50 ms transient load. Transient loads initiate the same feedback corrective responses as step loads, but lead to a substantial cessation of the feedback response 30 ms after the load is removed^48^. This suggests a spinal process can abruptly stop ongoing feedback corrections. Additional work is required to understand the relative contributions of spinal and supraspinal processes during co-contraction.

In short, we show that co-contraction improves corrective responses during postural and tracking tasks. While increasing muscle impedance does improve the perturbation response, the primary benefit of co-contraction appears to be the use of both agonist-antagonist muscles for feedback control. Our results, in combination with previous work, suggest that this dual agonist-antagonist control strategy is distinct from the common pattern of reciprocal inhibition between agonist and antagonists, and is triggered by active co-contraction. Importantly, the benefits of co-contraction are realized at relatively low levels of muscle activity, yielding performance improvements with low metabolic cost.

## Online Methods

All experiments were performed using a Kinarm Exoskeleton robot (Kinarm, Kingston, Ontario, Canada). Experiments were unimanual and participants used their dominant hand. Direct view of the upper limb was obstructed, and all visual feedback was provided using a virtual reality display aligned with the horizontal workspace. Upper limb kinematics were recorded by the robot and EMG was recorded for two elbow flexors (biceps and brachioradialis) and two elbow extensors (long and lateral heads of the triceps) using surface electrodes (Delsys Bagnoli, Delsys Inc., Natick, Massachusetts, USA). EMG recordings were mean centered, filtered (2-pass, 6th-order Butterworth, 20–250 Hz), rectified, and normalized^25^. At the beginning of both experiments, baseline EMG trials were performed. Subjects were required to maintain a static upper limb posture (45 degrees shoulder flexion and 90 degrees elbow flexion) while resisting torques of −5 to 5 Nm at the elbow. The mean muscle activity measured when resisting the 1 Nm loads in the baseline trials was used to normalize the EMG signals.

### Experiment 1 – Postural Perturbation

Twelve participants performed the first experiment. Participants were given visual feedback of their fingertip position and were presented with a 1 cm radius circular target at 45 degrees of shoulder flexion and 90 degrees of elbow flexion. The instructions for the task were to hold the fingertip in the target and to return to the target as quickly and accurately as possible when perturbed.

Trials consisted of three stages. During the first stage, participants brought their fingertip to the target and held it in the target for 2 seconds. At the completion of the first stage, visual feedback of the fingertip was removed, and the rest of the trial was performed using proprioceptive feedback. In the second stage, participants continued to hold their fingertip in the target, and a perturbation torque was applied by the robot to flex or extend the elbow at a random time between 2 and 4 seconds. The perturbation torque was a 5 Nm magnitude, 50 ms pulse with a 10 ms ramp up and ramp down. The third stage required participants to return their fingertip to the target and hold it inside the target. The trial was successful if the fingertip returned to the target within 500 ms of perturbation onset and stayed in the target for 1000 ms. Visual indication of trial success was provided (solid green circle – successful, solid red circle - unsuccessful). Feedback of the fingertip position was returned between trials.

Three types of trials were performed by the participants. During the first trial type, the robot applied a background flexion torque of 1, 3, or 5 Nm to activate the elbow extensors. Participants had to resist this background torque throughout all three stages of the trial. During these trials, the perturbation torque (flexion or extension) was applied in addition to the background torque. The second trial type was identical to the first, but the background torque was an extension torque that activated the flexors. In the first and second trial types the muscles pre-activated by the background load were referred to as the agonist muscles. The third trial type was performed without an applied background torque and with the participants actively co-contracting the muscles spanning the elbow to a target level of activity. The target levels of muscle activity were those required to resist the 1, 3, and 5 Nm loads during the baseline EMG trials. Real-time visual feedback of biceps activity, filtered using an exponential moving average, was given as a line that moved to the right as muscle activity increased. The participants had to hold their muscle activity at the target level, shown as a box spanning ±20% of the target, while keeping the fingertip at the target position during the first stage of the trial. EMG feedback was removed with fingertip feedback after the first stage. The three trial types resulted in three experimental loading conditions: (1) agonist muscles stretched by the perturbation, (2) agonist muscles shortened by the perturbation, and (3) muscles co-contracted.

Participants performed three groups of trials, one for each trial type. Each group consisted of one practice block of trials to adjust to the task, followed by three experimental blocks. Each trial type consisted of 30 trials completed in random order (3 background loads or 3 co-contraction levels x two perturbation directions x 5 repetitions).

Separate two-way repeated measures ANOVAs were performed for task success rate, peak elbow displacement, and return time using the loading condition (agonists stretched, agonists shortened, or muscles co-contracted) and level of muscle activity (1 Nm, 3 Nm, or 5 Nm equivalents) as factors. Significance values were adjusted for multiple comparisons using the Holm-Sidak method.

### Experiment 2 – Tracking Perturbation

Ten participants performed the second experiment. Participants were given visual feedback of their fingertip position and were presented with a 1 cm radius circular target that moved between 80 and 100 degrees of elbow flexion with shoulder flexion fixed at 45 degrees. The instructions for the task were to track the target with the fingertip as it moved across the display.

All trials started with the target at one end of the movement trajectory. Subjects moved their fingertip to the target and held it in the target for 2 seconds. The target moved to the other end of the trajectory and then returned to the start point at a constant angular speed of 4 deg/s. A perturbation was applied randomly between 87 and 93 degrees of elbow flexion during both the reach to the other end of the trajectory and the return to the starting point. The perturbation could apply an elbow torque in the same direction as the movement, a torque in the opposite direction of movement, or no torque. Each perturbation was equally likely, and perturbation torques were a 5 Nm magnitude, 50 ms pulse with a 10 ms ramp up and down. Participants performed trials with and without active antagonistic cocontraction of the muscles spanning the elbow. Unlike in the first experiment, no feedback of EMG activity was provided to the participants. They were simply instructed to perform the task with or without co-contraction.

Trial blocks consisted of nine trials in random order (there were nine possible perturbation combinations – 3 perturbation types for the initial reach x 3 perturbation types for the return movement). Participants performed two practice blocks, one with voluntary co-contraction and one without, to adjust to the task. Once adjusted, eight blocks of trials alternating between voluntary cocontraction and no co-contraction were performed. The endpoint of the trajectory that trials started from alternated every other block.

A two-way repeated measures ANOVA was performed for both the maximum displacement from the target and the return time to the target following the perturbation. Co-contraction (with or without) and the perturbation direction (same as or opposite to movement direction) were used as the factors. Significance values were adjusted for multiple comparisons using the Holm-Sidak method.

## Model

### Model Definition

We modeled the forearm and hand as a rigid link with a pin joint at one end (I). The link was controlled by two opposing torque actuators – one actuator for the elbow extensors (T_ext_) and one for the elbow flexors (T_flx_). The actuators could only produce positive torques, mimicking the ability of muscle to generate only tensile force. Stiffness (K) and damping (G) elements at the joint represented the passive impedance of the elbow joint from muscle and soft tissue. Equation 1 defines the model equation of motion and Equations 2 and 3 model the first order activation dynamics of muscle, where u is the control signal and τ is the time constant. Equation 4 defines the perturbation torque (T_pert_) as a step pulse.

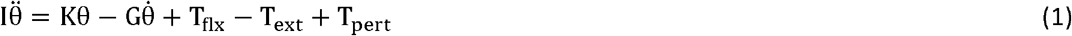

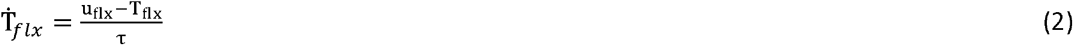

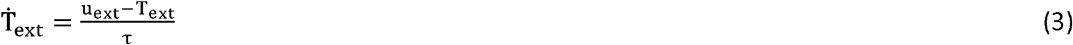

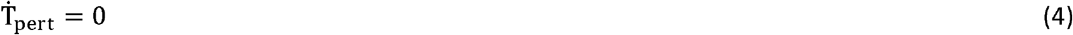

The model dynamics were discretized using an Euler method and a 10 ms time step. The discretized dynamics are defined in Equations 5–7. Note the addition of noise (ξ) to the state transition that affected only the applied torques. The additive noise was Gaussian with zero mean and variance of 1e-3.

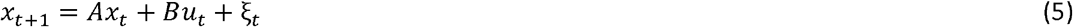

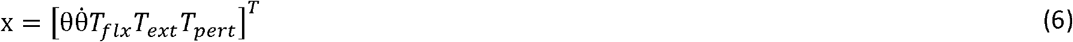

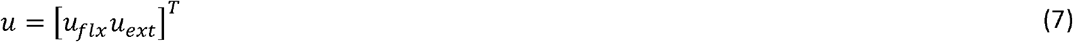

To account for feedback delays of h time steps, the discretized system was augmented to include state information from h previous time steps in Equations 8–12.

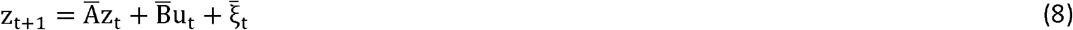

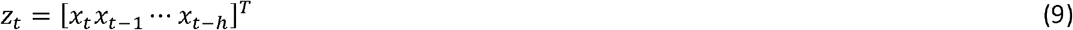

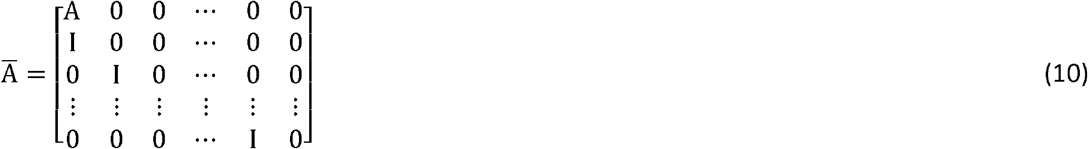

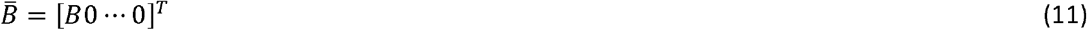

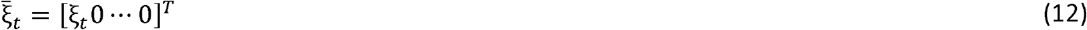

Delayed feedback was obtained from the augmented state as described in Equations 13 and 14, which simply define the feedback as the most delayed state (x_t-h_) plus additive noise. Additive noise was Gaussian with zero mean and variance of 1e-3.

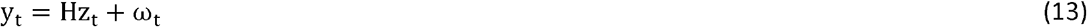

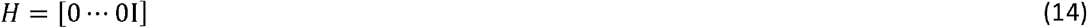

Linear-Quadratic-Gaussian control was used to derive an optimal feedback controller and optimal state estimator for the model. The controller feedback gains (L) were linear in the system state and the state estimator (S) was a Kalman filter. Equation 15 defines how the control at time t was calculated from the estimated state at time t. Equation 16 defines the update step for the estimated state, including the addition of Gaussian noise (η) with zero mean and variance 1e-3.

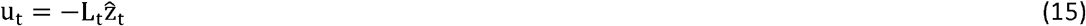

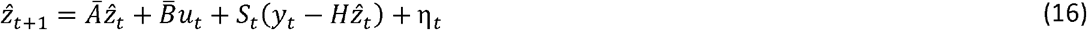

The stages of each simulation step are as follows:

1. Calculate the current estimated state (Equation 16).
2. Calculate the control from the current estimated state (Equation 15).
3. Advance the state to the next time step (Equation 8).

The system was initialized to a state of *x*_0_ = [00000]^*T*^ and a perturbation was applied by setting T_pert_ to 5 Nm for 5 time steps. Co-contraction was simulated by changing the minimum activation achievable by the actuators (i.e.: if simulating baseline activity of 1, then the minimum activation would be −1) and by changing the stiffness and damping at the joint.

### Model Parameters

The system inertia was estimated as I = 0.11 kg m^2^ based on average human anthropometry and the robot linkage inertia^19,49^. The time constant of the actuator dynamics was τ = 66 ms based on a first-order approximation of muscle dynamics^28^. The estimation of stiffness and damping were based on Force-length and force-velocity relationships of muscle described in literature^28,29,50^. The derivation process has been previously described^19^. The stiffness and damping values used for different levels of muscle activity are given in Table S3. Feedback was delayed by 6 time steps (h = 6), corresponding to long-latency sensorimotor delays^19,51^.

The cost minimized by the LQG controller is given in Equation 17.

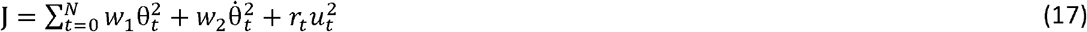

The cost prioritized minimizing position errors and the weighting parameters were adjusted to achieve return times of ~500 ms (w_1_ = 0.01, w_2_ = 1e-4, r = 1e-4).

## Acknowledgements

We thank Kim Moore, Sean Dukelow and Helen Bretzke for their technical support. This research was supported by NSERC Discovery grants to SHS, KD, MR and an ORF-RE grant to SHS and KD. SHS is also supported by a GSK Chair in Neuroscience.

## Author Contributions

All authors contributed to the conception and design of the experiments and model. CS performed the experiments, analyzed the data, developed the computational models, wrote the initial draft of the paper. All authors contributed to writing the manuscript.

## Tables (Online Methods)

**Table S1:**
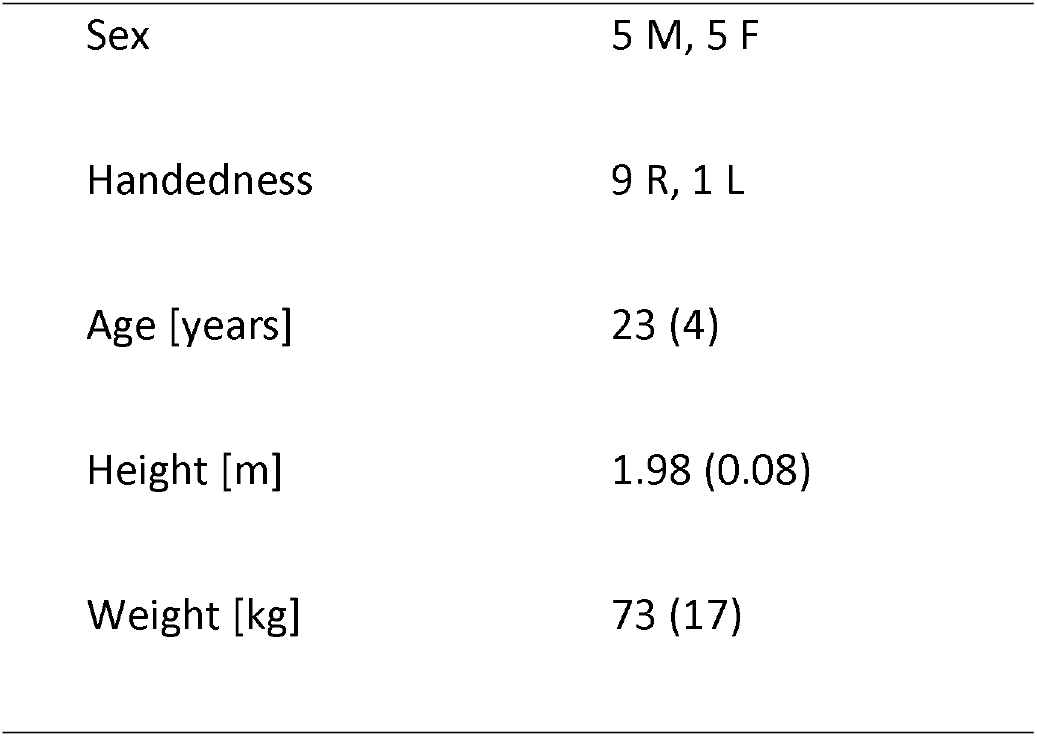
Experiment 1 subject demographics

**Table S2:**
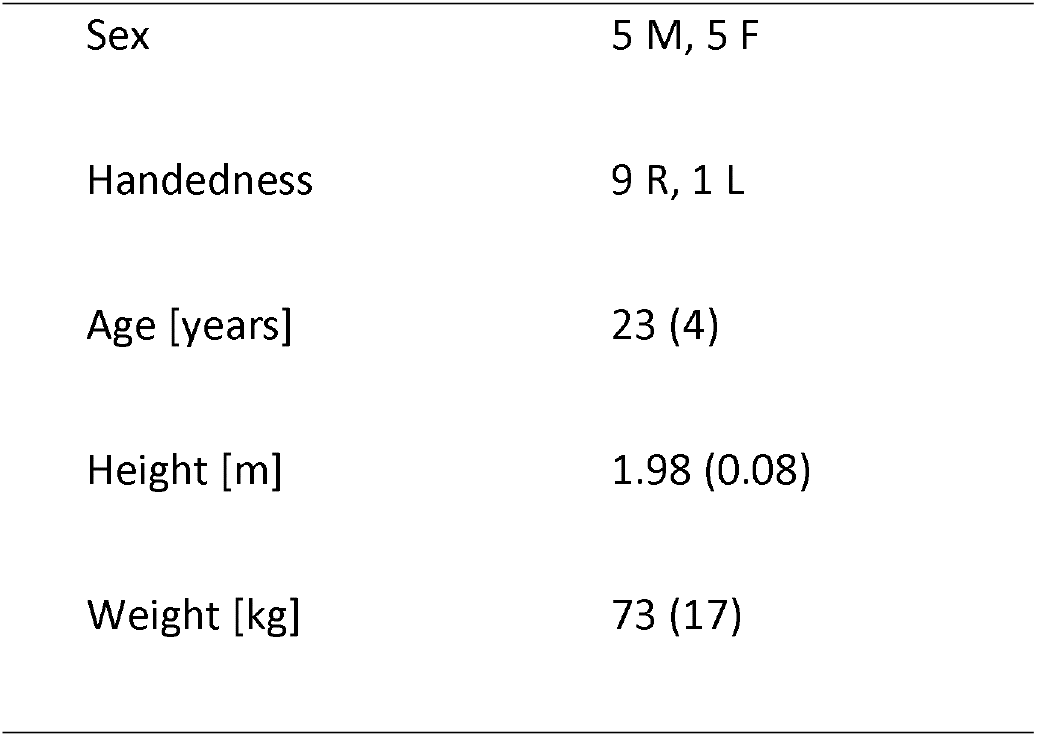
Experiment 2 subject demographics

**Table S3:** Stiffness and damping constants used in simulation

| Activity Level | Stiffness (K) | Damping (G) |
| --- | --- | --- |
| [Nm equivalent] | [Nm/rad] | [Nms/rad] |
| 1 | 1.55 | 0.18 |
| 3 | 3.28 | 0.45 |
| 5 | 4.40 | 0.73 |

## References

1. Hogan, N. Adaptive Control of Mechanical Impedance by Coactivation of Antagonist Muscles. IEEE Trans. Automat. Contr. 29, 681–690 (1984).

2. Reschechtko, S., Johansson, A. S. & Andrew Pruszynski, J. Maintaining arm control during self-triggered and unpredictable unloading perturbations. Eur. J. Neurosci. 50, 3531–3543 (2019).

3. Franklin, D. W., Osu, R., Burdet, E., Kawato, M. & Milner, T. E. Adaptation to Stable and Unstable Dynamics Achieved By Combined Impedance Control and Inverse Dynamics Model. J. Neurophysiol. 90, 3270–3282 (2003).

4. Milner, T. E. & Franklin, D. W. Impedance control and internal model use during the initial stage of adaptation to novel dynamics in humans. J. Physiol. 567, 651–664 (2005).

5. Osu, R. et al. Short- and Long-Term Changes in Joint Co-Contraction Associated With Motor Learning as Revealed From Surface EMG. J. Neurophysiol. 88, 991–1004 (2002).

6. Thoroughman, K. A. & Shadmehr, R. Electromyographic correlates of learning an internal model of reaching movements. J. Neurosci. 19, 8573–88 (1999).

7. Chambers, A. J. & Cham, R. Slip-related muscle activation patterns in the stance leg during walking. Gait Posture 25, 565–572 (2007).

8. Hogan, N. Impedance control: An approach to manipulation: Part I-theory. J. Dyn. Syst. Meas. Control. Trans. ASME 107, 1–7 (1985).

9. Franklin, D. W. & Wolpert, D. M. Computational Mechanisms of Sensorimotor Control. Neuron 72, 425–442 (2011).

10. Loeb, G. E., Brown, I. E. & Cheng, E. J. A hierarchical foundation for models of sensorimotor control. Exp. Brain Res. 126, 1–18 (1999).

11. Burdet, E. et al. A method for measuring endpoint stiffness during multi-joint arm movements. J. Biomech. 33, 1705–1709 (2000).

12. Gomi, H. & Osu, R. Task-dependent viscoelasticity of human multijoint arm and its spatial characteristics for interaction with environments. J. Neurosci. 18, 8965–8978 (1998).

13. Krutky, M. A., Trumbower, R. D. & Perreault, E. J. Influence of environmental stability on the regulation of end-point impedance during the maintenance of arm posture. J. Neurophysiol. 109, 1045–1054 (2013).

14. Mussa-Ivaldi, F. A., Hogan, N. & Bizzi, E. Neural, mechanical, and geometric factors subserving arm posture in humans. J. Neurosci. 5, 2732–43 (1985).

15. Perreault, E., Kirsch, R. & Crago, P. Effects of voluntary force generation on the elastic components of endpoint stiffness. Exp. Brain Res. 141, 312–323 (2001).

16. Tsuji, T., Morasso, P. G., Goto, K. & Ito, K. Human hand impedance characteristics during maintained posture. Biol. Cybern. 72, 475–485 (1995).

17. Matthews, P. B. C. The human stretch reflex and the motor cortex. Trends Neurosci. 14, 87–91 (1991).

18. Strick, P. L. Cerebellar involvement in volitional muscle responses to load changes. Cereb. Mot. Control man Long loop Mech. 4, 85–93 (1978).

19. Crevecoeur, F. & Scott, S. H. Beyond Muscles Stiffness: Importance of State-Estimation to Account for Very Fast Motor Corrections. PLOS Comput. Biol. 10, e1003869 (2014).

20. Scott, S. H. Apparatus for measuring and perturbing shoulder and elbow joint positions and torques during reaching. J. Neurosci. Methods 89, 119–127 (1999).

21. Bedingham, W. & Tatton, W. G. Dependence of EMG Responses Evoked by Imposed Wrist Displacements on Pre-existing Activity in the Stretched Muscles. Can. J. Neurol. Sci. / J. Can. des Sci. Neurol. 11, 272–280 (1984).

22. Marsden, C. D., Merton, P. A. & Morton, H. B. Servo action in the human thumb. J. Physiol. 257, 1–44 (1976).

23. Stein, R. B., Hunter, I. W., Lafontaine, S. R. & Jones, L. A. Analysis of short-latency reflexes in human elbow flexor muscles. J. Neurophysiol. 73, 1900–1911 (1995).

24. Verrier, M. C. Alterations in H reflex magnitude by variations in baseline EMG excitability. Electroencephalogr. Clin. Neurophysiol. 60, 492–499 (1985).

25. Pruszynski, J. A., Kurtzer, I., Lillicrap, T. P. & Scott, S. H. Temporal evolution of ‘automatic gain-scaling’. J. Neurophysiol. 102, 992–1003 (2009).

26. Hore, J. & Vilis, T. Loss of set in muscle responses to limb perturbations during cerebellar dysfunction. J. Neurophysiol. 51, 1137–48 (1984).

27. Lacquaniti, F. & Maioli, C. Anticipatory and reflex coactivation of antagonist muscles in catching. Brain Res. 406, 373–378 (1987).

28. Brown, I. E., Cheng, E. J. & Loeb, G. E. Measured and modeled properties of mammalian skeletal muscle. II. The effects of stimulus frequency on force-length and force-velocity relationships. J. Muscle Res. Cell Motil. 20, 627–643 (1999).

29. Li, W. & Todorov, E. Iterative linearization methods for approximately optimal control and estimation of non-linear stochastic system. Int.J. Control 80, 1439–1453 (2007).

30. Milner, T. E. Adaptation to destabilizing dynamics by means of muscle cocontraction. Exp. Brain Res. 143, 406–416 (2002).

31. Gribble, P. L., Mullin, L. I., Cothros, N. & Mattar, A. Role of Cocontraction in Arm Movement Accuracy. J. Neurophysiol. 89, 2396–2405 (2003).

32. Brown, I. E. & Loeb, G. E. A Reductionist Approach to Creating and Using Neuromusculoskeletal Models, in Biomechanics and Neural Control of Posture and Movement 148–163 (Springer New York, 2000). doi:10.1007/978-1-4612-2104-3_10

33. Joyce, G. C., Rack, P. M. H. & Westbury, D. R. The mechanical properties of cat soleus muscle during controlled lengthening and shortening movements. J. Physiol. 204, 461–474 (1969).

34. Rack, P. M. H. & Westbury, D. R. The short range stiffness of active mammalian muscle and its effect on mechanical properties. J. Physiol. 240, 331–350 (1974).

35. Morgan, D. L. Separation of active and passive components of short-range stiffness of muscle. Am. J. Physiol. Physiol. 232, 45–49 (1977).

36. Baldissera, F., Hultborn, H. & Illert, M. Integration in Spinal Neuronal Systems, in Comprehensive Physiology 509–595 (John Wiley & Sons, Inc., 2011). doi:10.1002/cphy.cp010212

37. Jankowska, E. Interneuronal relay in spinal pathways from proprioceptors. Prog. Neurobiol. 38, 335–378 (1992).

38. Lundberg, A. & Voorhoeve, P. Effects from the Pyramidal Tract on Spinal Reflex Arcs. Acta Physiol. Scand. 56, 201–219 (1962).

39. Crone, C. & Nielsen, J. Central control of disynaptic reciprocal inhibition in humans. Acta Physiol. Scand. 152, 351–363 (1994).

40. Tanaka, R. Reciprocal Ia inhibition during voluntary movements in man. Exp. Brain Res. 21, S125–S126 (1974).

41. Hultborn, H., Illert, M. & Santini, M. Convergence on Interneurones Mediating the Reciprocal Ia Inhibition of Motoneurones III. Effects from supraspinal pathways. Acta Physiol. Scand. 96, 368–391 (1976).

42. Vallbo, A. B. Human Muscle Spindle Discharge during Isometric Voluntary Contractions. Amplitude Relations between Spindle Frequency and Torque. Acta Physiol. Scand. 90, 319–336 (1974).

43. Nielsen, J. & Kagamihara, Y. The regulation of disynaptic reciprocal Ia inhibition during co-contraction of antagonistic muscles in man. J. Physiol. 456, 373–391 (1992).

44. Nielsen, J. & Kagamihara, Y. The regulation of presynaptic inhibition during co-contraction of antagonistic muscles in man. J. Physiol. 464, 575–593 (1993).

45. Hultborn, H., Jankowska, E. & Lindström, S. Relative contribution from different nerves to recurrent depression of Ia IPSPs in motoneurones. J. Physiol. 215, 637–664 (1971).

46. Fetz, E. E. & Cheney, P. D. Functional relations between primate motor cortex cells and muscles: fixed and flexible. Ciba Found. Symp. 132, 98–117 (1987).

47. Scott, S. H. Putting Sensory Back into Voluntary Motor Control. in 3–7 (Springer, Singapore, 2016). doi:10.1007/978-981-10-0207-6_1

48. Kurtzer, I., Pruszynski, J. A. & Scott, S. H. Long-Latency and Voluntary Responses to an Arm Displacement Can Be Rapidly Attenuated By Perturbation Offset. J. Neurophysiol. 103, 3195–3204 (2010).

49. Winter, D. A. Biomechanics and Motor Control of Human Movement: Anthropometry. (Wiley, 2004).

50. Scott, S. H., Brown, I. E. & Loeb, G. E. Mechanics of feline soleus: I. Effect of fascicle length and velocity on force output. J. Muscle Res. Cell Motil. 17, 207–219 (1996).

51. Scott, S. H. The computational and neural basis of voluntary motor control and planning. Trends in Cognitive Sciences 16, 541–549 (2012).

