## Supplemental Figures for "Co-contraction uses dual control of agonist-antagonist muscles to improve motor performance"

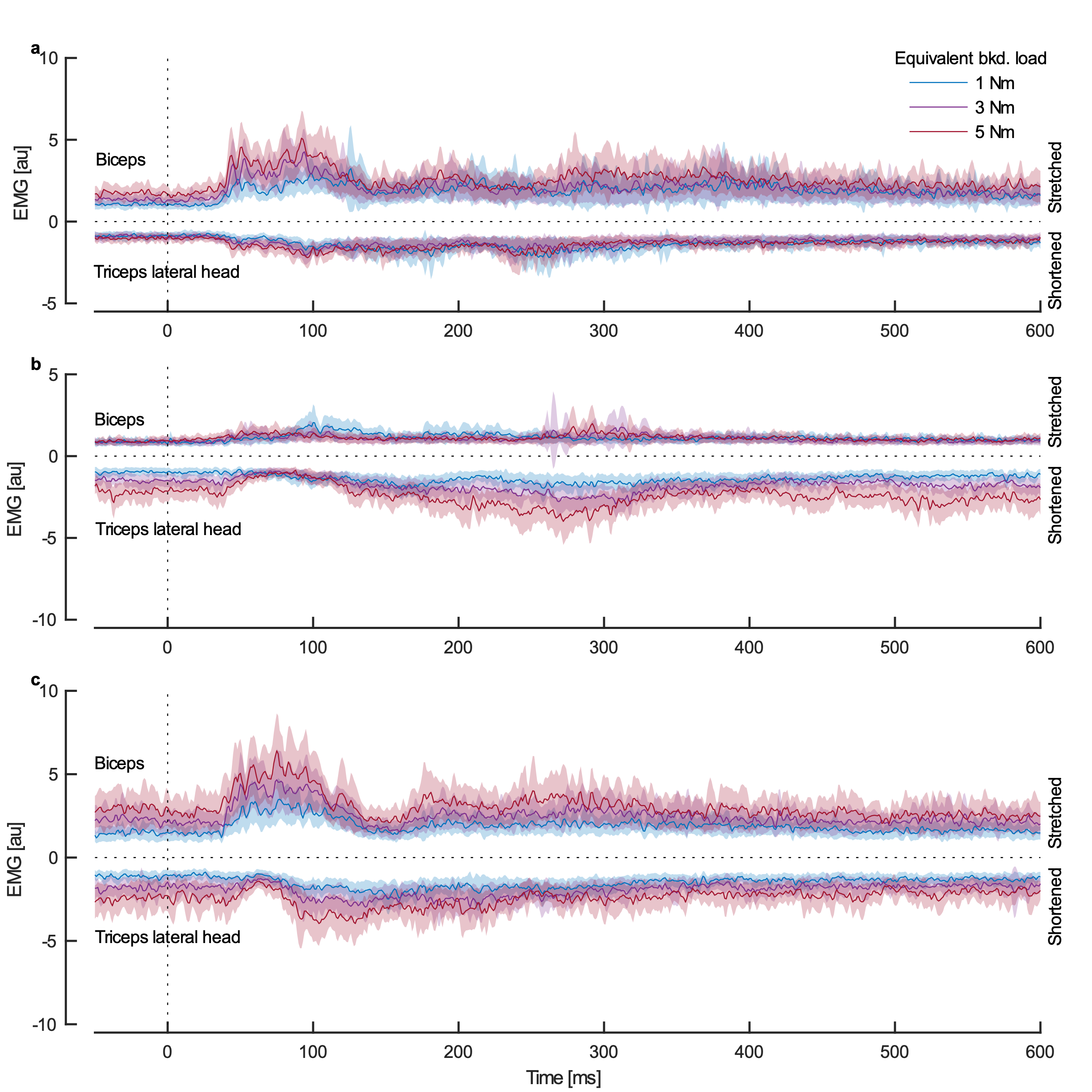

Supplemental Figure 1: Activity of the triceps lateral head and biceps following mechanical perturbations that extend the elbow. (a) Stretched biceps pre-activated by background load. (b) Shortened triceps pre-activated by background load. (c) Muscles co-contracted. Increases in activity of the stretched biceps are plotted in the positive direction and increases in activity of the shortened triceps are plotted in the negative direction. Background muscle activity level is indicated by line color and perturbation onset is at 0 ms. Group results (mean and standard error) are shown. Note that while the stretched biceps does not display distinct short and long latency bursts, its response is still attenuated when unloaded (b) but not when co-contracting (c), demonstrating dual agonist-antagonist control.

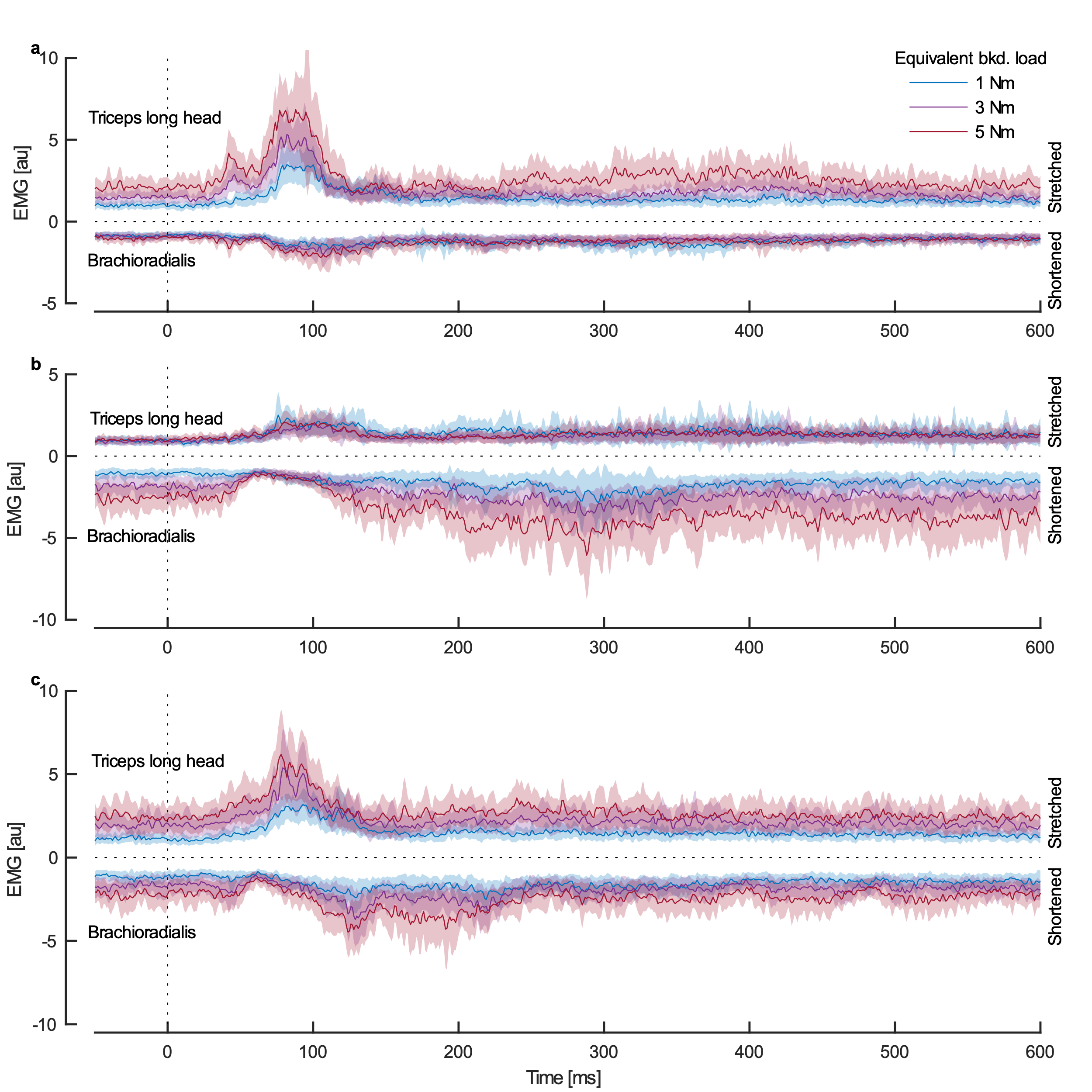

Supplemental Figure 2: Activity of the triceps long head and brachioradialis following mechanical perturbations that flex the elbow. (a) Stretched triceps pre-activated by background load. (b) Shortened brachioradialis pre-activated by background load. (c) Muscles co-contracted. Increases in activity of the stretched triceps are plotted in the positive direction and increases in activity of the shortened brachioradialis are plotted in the negative direction. Background muscle activity level is indicated by line color and perturbation onset is at 0 ms. Group results (mean and standard error) are shown.

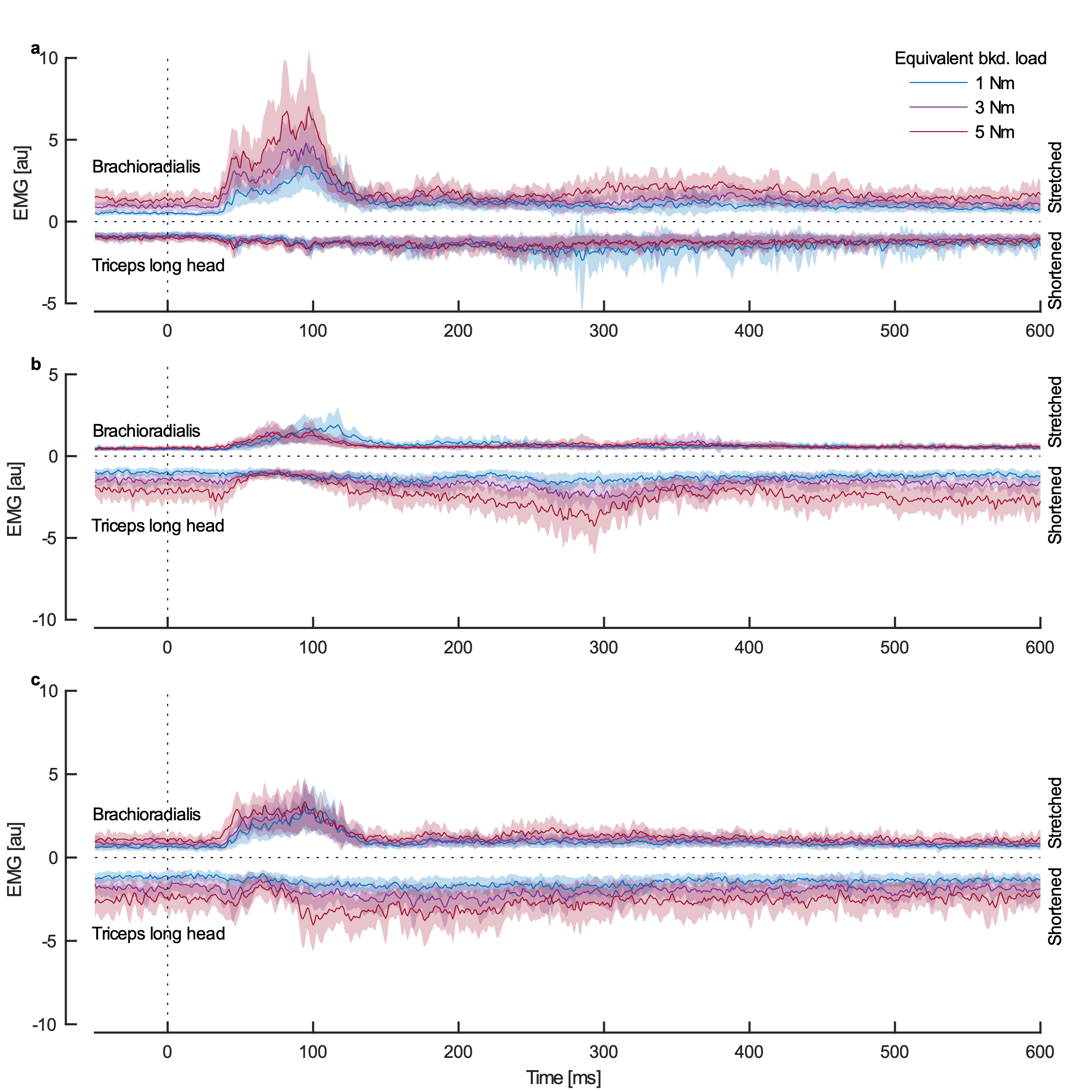

Supplemental Figure 3: Activity of the triceps long head and brachioradialis following mechanical perturbations that extend the elbow. (a) Stretched brachioradialis pre-activated by background load. (b) Shortened triceps pre-activated by background load. (c) Muscles co-contracted. Increases in activity of the stretched brachioradialis are plotted in the positive direction and increases in activity of the shortened triceps are plotted in the negative direction. Background muscle activity level is indicated by line color and perturbation onset is at 0 ms. Group results (mean and standard error) are shown. Note that while the attenuation of the stretched brachioradialis is not completely eliminated when co-contracting (c), it is still more involved in the response than when only the triceps is pre-activated (b), supporting dual agonist-antagonist control.
